# Adult Mouse Leptomeninges Exhibit Regional and Age-related Cellular Heterogeneity Implicating Mental Disorders

**DOI:** 10.1101/2023.09.10.557097

**Authors:** Christina A. Allen, Susan K. Goderie, Mo Liu, Thomas R. Kiehl, Farhad Farjood, Yue Wang, Nathan C. Boles, Sally Temple

## Abstract

The leptomeninges envelop the central nervous system (CNS) and contribute to cerebrospinal fluid (CSF) production and homeostasis. We analyzed the meninges overlying the anterior or posterior forebrain in the adult mouse by single nuclear RNA-sequencing (snucRNA-seq). This revealed regional differences in fibroblast and endothelial cell composition and gene expression. Surprisingly, these non-neuronal cells co-expressed genes implicated in neural functions. The regional differences changed with aging, from 3 to 18 months. Cytokine analysis revealed specific soluble factor production from anterior vs posterior meninges that also altered with age. Secreted factors from the leptomeninges from different regions and ages differentially impacted the survival of anterior or posterior cortical neuronal subsets, neuron morphology, and glia proliferation. These findings suggest that meningeal dysfunction in different brain regions could contribute to specific neural pathologies. The disease-associations of meningeal cell genes differentially expressed with region and age were significantly enriched for mental and substance abuse disorders.

## Introduction

The leptomeninges surround the brain and spinal cord.^1 2,3^ They maintain and protect the central nervous system (CNS), performing multiple functions including acting as a physical barrier, contributing to immune surveillance, and regulating cerebrospinal fluid production and composition. The leptomeninges are composed of a diverse array of cells, including fibroblasts, resident and transient macrophages, blood vessel endothelial cells, mural cells, and immune cells. The outer dura mater is the periosteal layer that provides structural support to the brain and CNS lymphatic system. The inner arachnoid and pia mater layers, collectively referred to as the leptomeninges, surround the subarachnoid space and blanket the CNS, penetrating the perivascular spaces in the brain. Leptomeningeal-to-brain interactions support brain functions under steady-state, inflamed, and diseased conditions through production of cytokines and growth factors.^4^

The leptomeninges play important roles during development by signaling to underlying neural cells. For example, they secrete retinoic acid to stimulate neuron production during corticogenesis^5,6^ and they secrete the ligand CXCL12 to guide migration of CXCR4-expressing cells such as Cajal-Retzius neurons.^7^ A recent single cell RNA sequencing study indicates the developing leptomeninges dissected from different embryonic mouse brain regions have different fibroblast composition.^8^ Much less is known about the trophic support and signaling provided by the leptomeninges to underlying brain tissue in adulthood. Prior studies indicate that they continue to provide retinoic acid to regulate germinal zones in the hippocampus,^9^ but it is not known if the adult leptomeninges are regionally specified, producing diverse factors specialized to support subjacent neural cells. Considering the vast area and diverse CNS regions covered by the leptomeninges, we questioned the generally held assumption that they function as a uniform supportive structure.

Here, we demonstrate that in the adult mouse, the leptomeninges dissected from anterior versus posterior forebrain regions exhibit distinct cellular composition by single nuclear sequencing (snucRNA-seq). Young leptomeninges included three clusters of endothelial cells and three clusters of fibroblasts that show clear regional separation. With age, clusters retain regional differences but differences between cell subtypes are less defined. A fibroblast cluster *Fibro2* and an endothelial cluster, and *Endo3*, were significantly reduced with aging. Using secreted protein analysis, we demonstrated specific cytokine enrichment in anterior and posterior leptomeninges and significant changes with age. We then used a co-culture assay to show that the regional and age-associated differences in factors produced by the leptomeninges impact the survival and growth of anterior versus posterior cortical neurons and glia. These results demonstrate that the leptomeninges are not uniform, but rather provide support that is regionally specified and that age-associated changes in leptomeningeal function, such as enhanced gliogenic factor production and increased inflammatory factor production, could impact brain aging and health in a region-related manner. Analysis of differentially expressed genes in anterior and posterior forebrain leptomeninges and how these change with age highlighted associations with mental illnesses including schizophrenia and bipolar disease.

## Results

### Single cell sequencing reveals regional and age-related differences in endothelial cell and fibroblast populations

The leptomeninges consist of a highly heterogeneous population of resident and transient cell types. To distinguish the different populations and molecular identities of leptomeningeal cells in different regions and investigate changes with age, we performed snucRNA-seq with the iCELL8 chip-based system. The leptomeninges were dissected from the forebrain anterior region overlying the frontal and parietal-temporal cortical lobes and the posterior region overlying the occipital cortex, subicular cortex, entorhinal cortex and hippocampus (Fig. S1A). snucRNA-seq data was mapped with STAR^10^ and gene counts, including exons and introns, were used for downstream analysis. After QC, cells had library sizes of at least 30,000 reads (median of ∼60,000) and over 1000 genes per cell (median of 3750); we studied 722 cells from 3-month old anterior leptomeninges, 1179 from 3-month posterior leptomeninges, and 640 each from 18-month old anterior and posterior 18-month old leptomeninges. Snuc-RNA expression was analyzed with the Seurat package (v.4.1.1) in R. Initial processing through the Seurat pipeline demonstrated a primary driver of variance was the anterior or posterior region indicating important regional differences (Fig. 1A-1H). The vast majority of the recovered cells were in endothelial or fibroblast categories, which we focused on in subsequent analysis.

**Figure 1.**
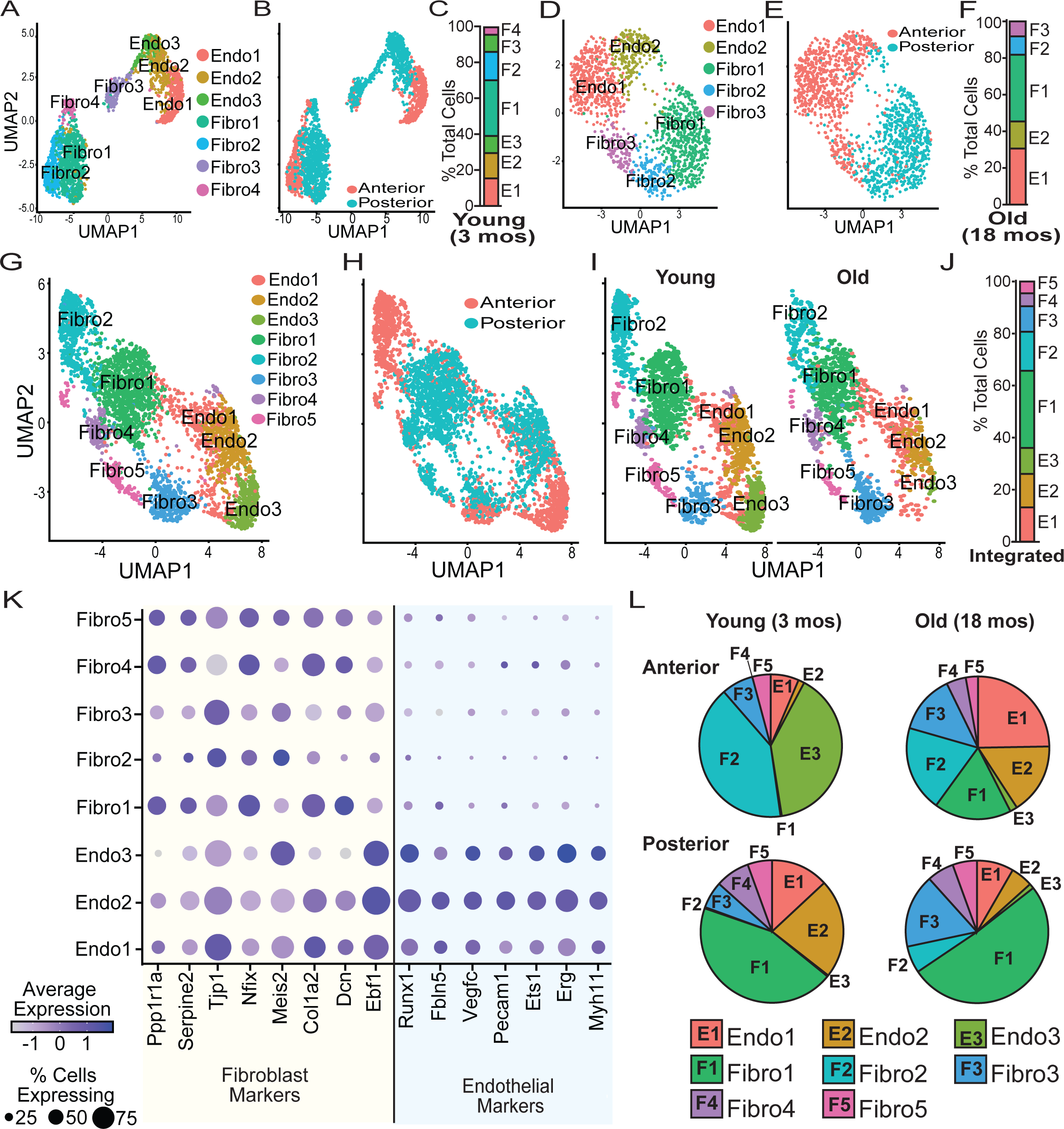
Single nuclei RNA (snRNA-seq) sequencing identified significant differences in the cellular composition of leptomeninges between regions and ages. Leptomeninges was dissected from the anterior and posterior of young (3mo) and old (18mo) mice. (A-B) Young leptomeninges demonstrated a clear distribution of cells based on region of isolation and cell identity with 4 fibroblastic and 3 endothelial subpopulations. (C) In the young leptomeninges the fibroblast populations composed approximately 60% of the total cells with the remainder being endothelial cell subpopulations. (E-F) Old leptomeninges also demonstrated a distribution of cells based on region of isolation and cell identity with 3 fibroblastic and 2 endothelial subpopulations. (F) In the old leptomeninges, roughly 55% of the cells are fibroblasts with the remainder falling in the endothelial group. (G-H) The integrated dataset retained the strong biases in the distribution of cells by region with a total of three 3 endothelial and 5 fibroblastic subpopulations. (I) Cells for each group showed representation across ages. (J) The composition of the total cells, approximately 60/40 for fibroblasts/endothelial cells. (K) Dotplot of fibroblast and endothelial markers used to assign cluster identity. (L) Distribution of cell subpopulations split by region and age.

In the young leptomeninges (3-months), we identified seven cell major types: four fibroblast populations and three endothelial cell populations. A clear delineation between young anterior and young posterior cell populations was seen in the UMAP plots (Fig. 1A-1C). In the old leptomeninges (18-months), we identified two endothelial populations, and three fibroblastic populations (Fig. 1D-1F). We then integrated the data from both ages using canonical correlation analysis (CCA) (Fig. 1G-1J). Regionality remained a strong driver of variance in the integrated dataset. The integrated data revealed three endothelial cell populations but now five fibroblastic populations. The endothelial cell populations expressed markers including *Pecam1* (CD31), *Erg*, and *Ets1*, and the fibroblast populations expressed markers including *Tjp1*, *Col1a2*, and *Dcn* (Fig. 1K).

We noted significant differences in the different cell populations in each region and with aging (Fig. 1L). In young anterior leptomeninges, nearly half of recovered cells were endothelial cells, whereas in the young posterior leptomeninges only one third were endothelial cells. In the young anterior leptomeninges, the *Endo3* and *Fibro2* expressing subpopulations formed the majority of cells detected. However, both these populations were much reduced with aging, with the *Endo3* population being nearly absent in 18-month-old anterior leptomeninges. Interestingly, the cellular composition of the aged anterior leptomeninges was closer to the composition of the young posterior leptomeninges than the young anterior leptomeninges. The young posterior leptomeninges were composed largely of *Fibro1, Endo1* and *Endo2* populations. Some changes occurred in the cellular composition of the posterior leptomeninges with age such as the expansion of the *Fibro1* population and reductions in endothelial cell subpopulations, however the most drastic changes were observed in the anterior leptomeninges with age.

### Neural specialization and disease association in endothelial and fibroblast gene expression

Following, we examined differential gene expression in meningeal fibroblasts and endothelial cells between brain regions and ages and between cell types using the Wilcoxon rank sum test (Tables S1-S2). We then performed gene ontology (GO) enrichment analysis using the SCREP package^11^ semantic similarity analysis. In the age and region enrichment (Fig. 2A, Tables S3), neurogenesis-associated genes were highly enriched in the young ages, especially the posterior cells. All regions and ages were enriched for genes involved with cell morphogenesis. The posterior leptomeninges with aging were enriched for genes associated with programmed cell death. When examining the enrichments based on cell types, many of the same parent categories that were enriched by region and age were also highly enriched (Fig. 2B, Tables S4). The *Endo3* population had the lowest number of enriched categories (245) and the *Fibro1* population had the greatest number (1504). *Endo3* was the primary endothelial cell population in the young anterior leptomeninges, but this population is essentially lost with age. Nearly 20% of the enriched categories for *Endo3* fall under “cell projection organization” and “ion transport”. When compared to the other endothelial subpopulations, *Endo3* was uniquely enriched for categories dealing with “behavior”, “glycosylation”, “establishment of cell polarity”, and “developmental growth. The mouse brain continues to mature after birth^12^ and these unique categories may indicate a role for this cell population in postnatal brain development that is no longer needed when the brain is fully mature, leading to the loss and replacement of this cell population.

**Figure 2.**
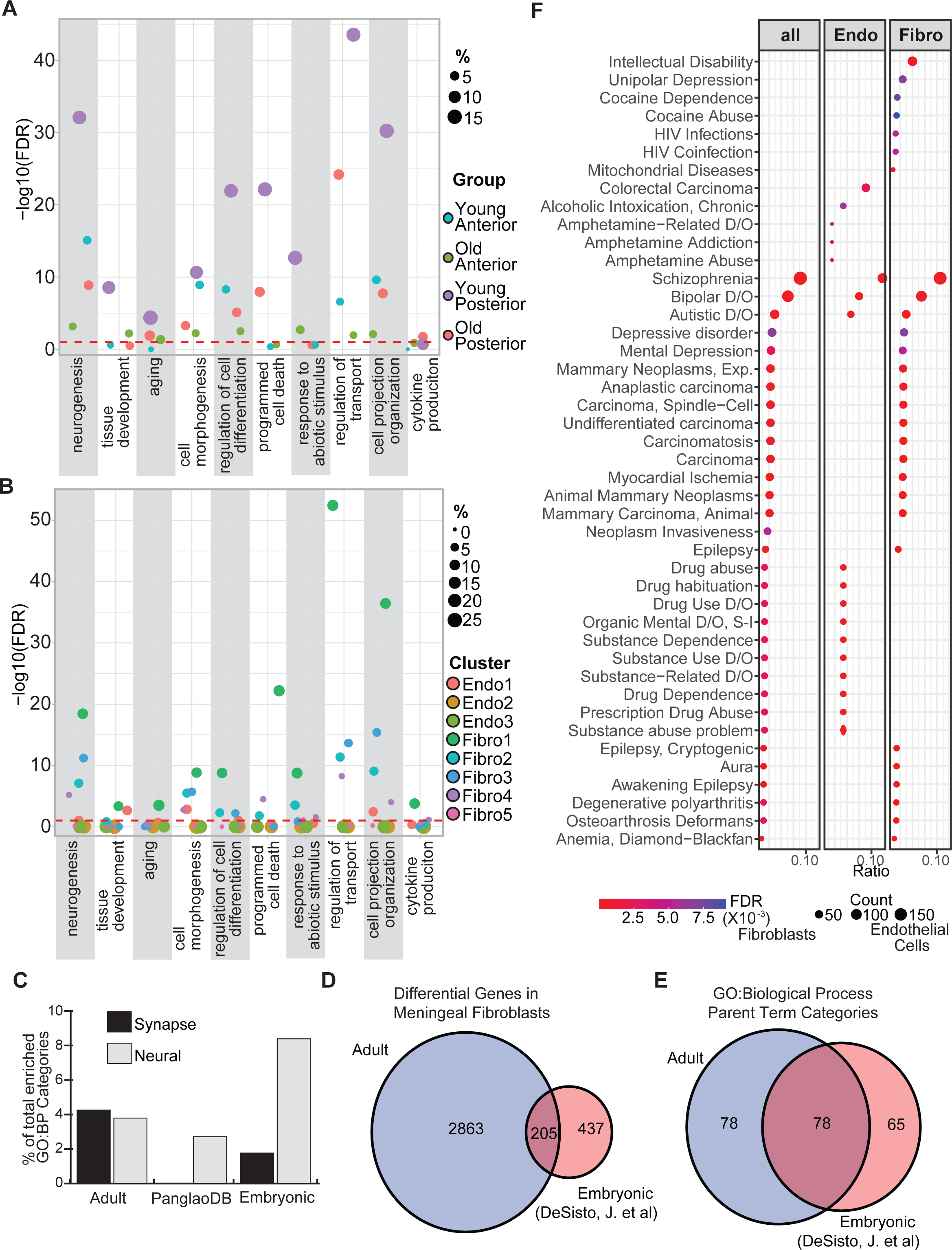
Enrichment analysis of snRNA-seq data from the leptomeninges. Marker genes were identified based on age and region (Table S1) or subpopulation (Table S2). Using the SCREP package, gene ontology enrichment followed by semantic similarity analysis was carried out for the markers based on age and region (A) or subpopulation (B) (Tables S3-S4). (C) Comparison of synapse and neural enriched gene ontology biological processes (GO:BP) categories in adult meningeal fibroblasts, embryonic meningeal fibroblasts and the PanglaoDB fibroblast markers. (D,E) Overlap of the adult meningeal fibroblast dataset and the embryonic meningeal fibroblast datasets comparing (D) differentially expressed (DEX) genes and (E) the GO:BP. (F) Disease enrichment was performed with the disgenet2r package. Markers tested were the complete intersections of Table S1 and Table S2 and the intersections based on dividing Table S2 into endothelial or fibroblastic subpopulations.

The *Fibro1* subpopulation was the primary fibroblastic cell in the posterior leptomeninges and the old anterior leptomeninges. Roughly 10% of its overall enriched categories fall under the parent terms “homeostatic process”, “ion transport” and “intracellular process”. The *Fibro2* population, which is a large proportion of cells in young anterior leptomeninges that decreases by ∼50% with age, shares most of its enriched categories with the *Fibro1* population, such as “morphogenesis”, “neurogenesis”, and “cell cycle”. However, examination of the child terms reveal differences between these two fibroblast populations. For example, under the parent category “cell cycle”, the child terms that deal with positive cell cycle regulation in the *Fibro1* population are absent in the *Fibro2* population. Among the fibroblast subpopulations, *Fibro2* has some exclusively enriched parent terms including “demethylation”, “maintenance of cell number” and developmental categories such as “cranial skeletal system development”. As with the *Endo3* subpopulation, the *Fibro2* cells may be playing a role in brain maturation. The loss of *Fibro2* cells with age is compensated by increases in *Fibro1* and *Fibro4* cells. Both of these subpopulations express APOE, a gene implicated in neurodegeneration that may have a role in shrinkage of the meningeal lymphatic system.^13^ Hence, changes in the leptomeningeal cell populations may contribute to neurodegenerative diseases such as Alzheimer’s disease, along with recognized changes in other meningeal functions such as lymphatic drainage.^14^

A surprising result from this enrichment analysis was the abundance of synapse-related pathways enriched in fibroblast populations. At least 3.6% (*Fibro 2*) and up to 9.3% (*Fibro3*) of enriched categories were related to synapse biology. To test if this was a general property of fibroblasts, we downloaded markers associated with fibroblasts in PanglaoDB (a compendium of single cell datasets across ∼250 tissues and ∼1400 samples)^15^ and performed enrichment testing (Fig. 2C). The PangalaoDB general fibroblast markers (177) did not show enrichment for categories associated with synapses, although a few nervous system related pathways (27/994) were found such as “neuron death” and “neuron migration”. We then compared our adult snucRNA-seq data to a previously published embryonic meningeal fibroblast single cell RNA-seq dataset.^8^ Approximately half of the embryonic fibro-meningeal marker genes overlapped with our adult leptomeningeal fibroblast markers (Fig. 2D). Following enrichment analysis based on the embryonic fibroblast subpopulation markers using the SCREP package, half of the enriched GO parent terms from the adult dataset were also in the embryonic dataset (Fig. 2E). Looking specifically at synapse-related GO categories, four of the eleven embryonic fibroblast subpopulations had enrichments for at least one synapse term but none showed the substantial synapse-related enrichments seen in the adult populations. As a further comparison, we examined the adult, the PanglaoDB, and the embryonic markers for enrichment of terms with the keyword “neural”. All three demonstrated enriched categories with this keyword (Fig. 2C), with the embryonic markers showing double the enriched terms compared to the adult or PanglaoDB markers. These results suggest a previously uncharacterized function for fibroblasts of the leptomeninges, and particularly the adult leptomeninges, in regulating synapse activity.

We next investigated the potential impact of the changes in gene expression between region, age, and cell type on disease. We intersected the differentially expressed genes identified between region and age (Table S1) with the genes associated with cell type (Table S2) and generated three gene lists:1) all genes in the intersection, 2) intersected genes from the fibroblast populations, 3) intersected genes from the endothelial cell populations. This identified the genes in endothelial cells and fibroblasts that were changed in anterior versus posterior leptomeninges and/or with age. We utilized the disgenet2r package^16^ to look at disease enrichment utilizing these three gene lists. Unexpectedly, enriched diseases were divided broadly into three categories: mental disorders, substance abuse disorders, and nervous system diseases such as epilepsy. Genes that were associated with age, region, and fibroblasts were enriched across all three of these major disease categories. In contrast, genes associated with region, age, and endothelial cell populations were primarily affiliated with substance abuse disorders. Genes associated with schizophrenia were highly enriched across the cell types. Together these data reveal a previously unrecognized regional identity of the meninges that is impacted by aging with potential repercussions across several neural disorders, including schizophrenia, substance abuse, and epilepsy.

### Identifying functional differences in neural support of factors from anterior and posterior leptomeninges

To assess the ability of the anterior or posterior leptomeninges to support underlying neural cells, we utilized homo- and heterotypic co-culture with embryonic day 17 (E17) anterior versus posterior cerebral cortical cells (Fig. 3A, S1A, S1B). E17 mouse cortical cells were chosen to provide a read-out of the meningeal support because fetal neurons survive better in culture than adult neurons and most cortical subtypes are born by E17, enabling us to assess survival of different cortical layer neurons as well as responses of glial populations.

**Figure 3.**
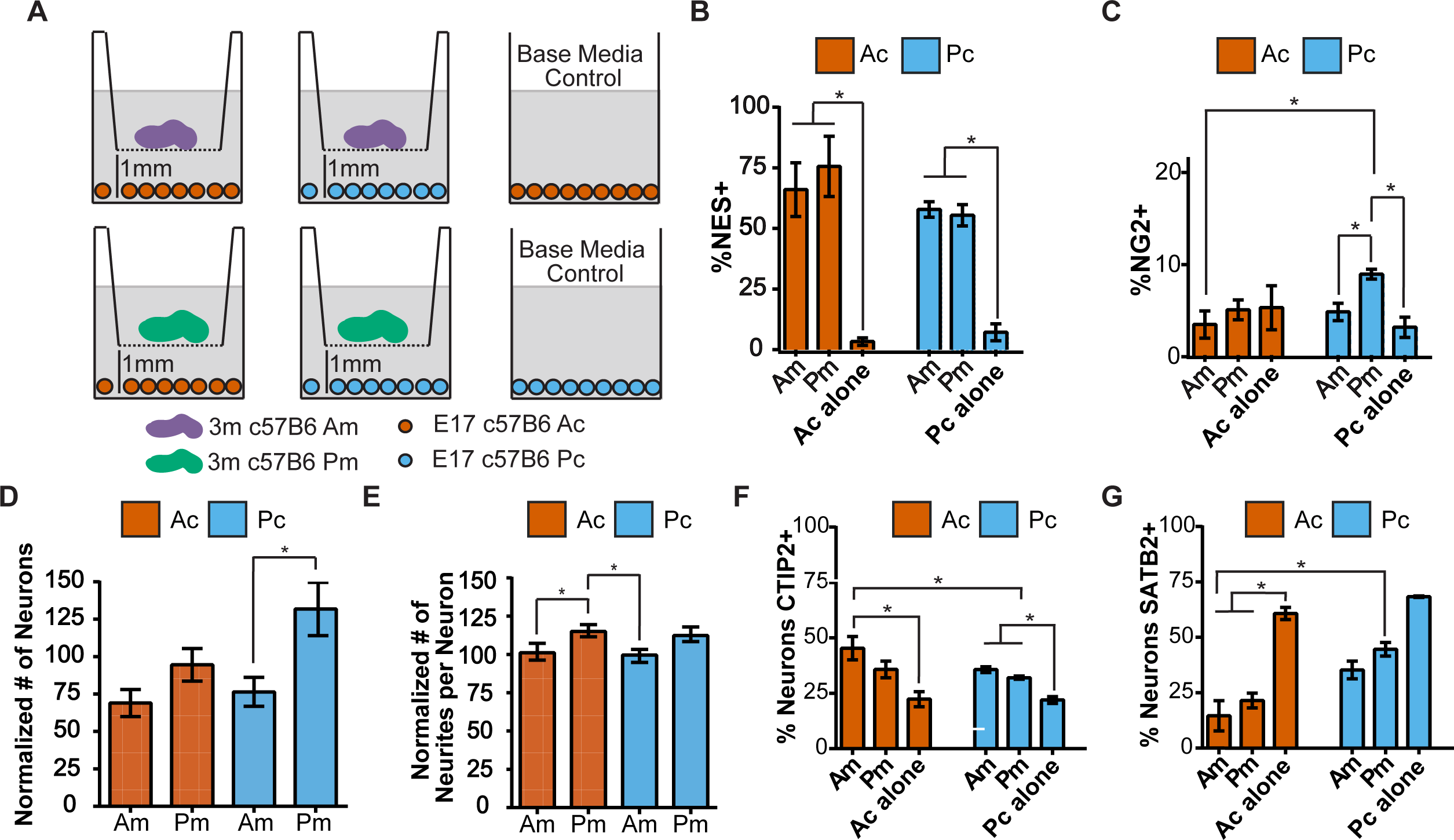
Anterior and posterior leptomeninges have different abilities to support growth of anterior and posterior cortical cultures. (A) Schematic of experimental design used to assess homo- and heterotypic combinations of anterior (Am, purple) and posterior (Pm, green) leptomeninges in transwells co-cultured with anterior (Ac, orange) or posterior (Pc, light blue) cortical cells. Control: cortical cultures grown without leptomeninges; (B-G) Cells were counted in five 10x image frames per experiment: (B) % of NES+ progenitor cells; (C) % of NG2+ cells; (D) The number of TuJ1+ cells (neurons) with Am or Pm co-culture normalized to the number of TuJ1+ cells in Ac or Pc alone control conditions. (E) The number of neurites per neuron in Am or Pm co-culture normalized to the average neurites per neuron in Ac or Pc alone control conditions; (F) % of neurons that are CTIP2^+^; (G) % of neurons that are SATB2^+^. Unpaired Student’s t-test was used to determine statistical significance(*=p<0.05). Error bars represent mean ± SEM.

Prior studies have demonstrated that the young leptomeninges produce factors that affect nearby progenitor cells at the pial surface.^6,17,18^ We found that posterior leptomeninges produced significantly more cell growth from co-cultured anterior and posterior cortex compared to anterior leptomeninges (Fig. S1C, S1D). After 4 days, immunostaining showed an expansion of Nestin (NES)^+^ neural progenitor cells (Fig. 3B, S2A-S2D). Anterior leptomeninges promoted significantly more NES+GFAP+ cells, presumed early astrocytes, from anterior cortical cells (Fig. S2B) although no differences in mature astrocytes (NES-GFAP+) were observed (Fig. S2C). In contrast, posterior leptomeninges co-culture produced significantly more NG2^+^, putative oligodendrocyte lineage cells from posterior cortical cells (Fig. 3C). These data demonstrate that the adult leptomeninges secrete paracrine factors that have powerful effects on glial cells and precursors that are regionally distinct, with anterior leptomeninges promoting expansion of astrocyte lineage cells in the anterior cortex and posterior leptomeninges promoting expansion of oligodendrocyte precursors in the posterior cortex.

We then investigated the impact of the leptomeninges on cerebral cortical neuron survival. Anterior leptomeninges co-culture reduced anterior and posterior neuron number compared to control, while posterior leptomeninges co-culture resulted in approximately 25% more posterior neurons compared to control (Fig. 3D, S2E). In addition, anterior leptomeningeal factors caused a reduction in anterior but not posterior cortical neurite length while posterior leptomeninges promoted more and longer neurites in anterior cortical co-cultures than did anterior leptomeninges (Fig. 3E, S2F-G). Interestingly, leptomeningeal co-culture resulted in more CTIP2+ (subcortical projection) neurons and fewer SATB2+ (callosal projection) neurons compared to cultures without leptomeninges. The anterior leptomeninges supported more CTIP2+ neurons in anterior cortical cultures than control (Ac alone), and the posterior leptomeninges supported more SATB2+ neurons in posterior cortical cultures than seen in anterior co-culture (Ac-Am) (Fig. 3F-G). Hence, the leptomeninges from different cortical regions release factors that can differentially affect the survival and morphology of subtypes of neurons in the underlying cortex.

### Aged leptomeninges show reduced support of glial progenitors and neurons

Histological studies have shown that the leptomeninges thicken and calcify with age.^19^ However, the impact of these changes on the ability to support subjacent neural tissue has not been assessed. To examine this, we utilized the same co-culture paradigm described above, but only in homotypic combination with anterior or posterior E17 cortical cells (Fig. 4A). Notably, we observed a distinct difference in cell survival: posterior leptomeninges increased Caspase 3 expression, a marker of apoptosis, in posterior cortical cells in an age-dependent manner, an effect not seen with anterior leptomeninges coculture (Fig. 4B). With age, leptomeninges showed decreased growth in the cortical cultures, an effect more prominent at 18-compared to 22-months (Fig. 4C), corroborating prior studies of aging forebrain progenitor cells that show less proliferation at 18-compared to 22-months.^20^ 18-month-old leptomeninges coculture resulted in fewer NES^+^ progenitor cells and more NES-GFAP+ astrocytes than co cultured with either 3- or 22-month leptomeninges (Fig. 4D-E). In contrast to these non-monotonic trends, co-cultures with aging posterior leptomeninges resulted in steadily fewer NG2^+^ cells through 22 months, an effect not observed in co-cultures with anterior leptomeninges (Fig. 4F).

**Figure 4.**
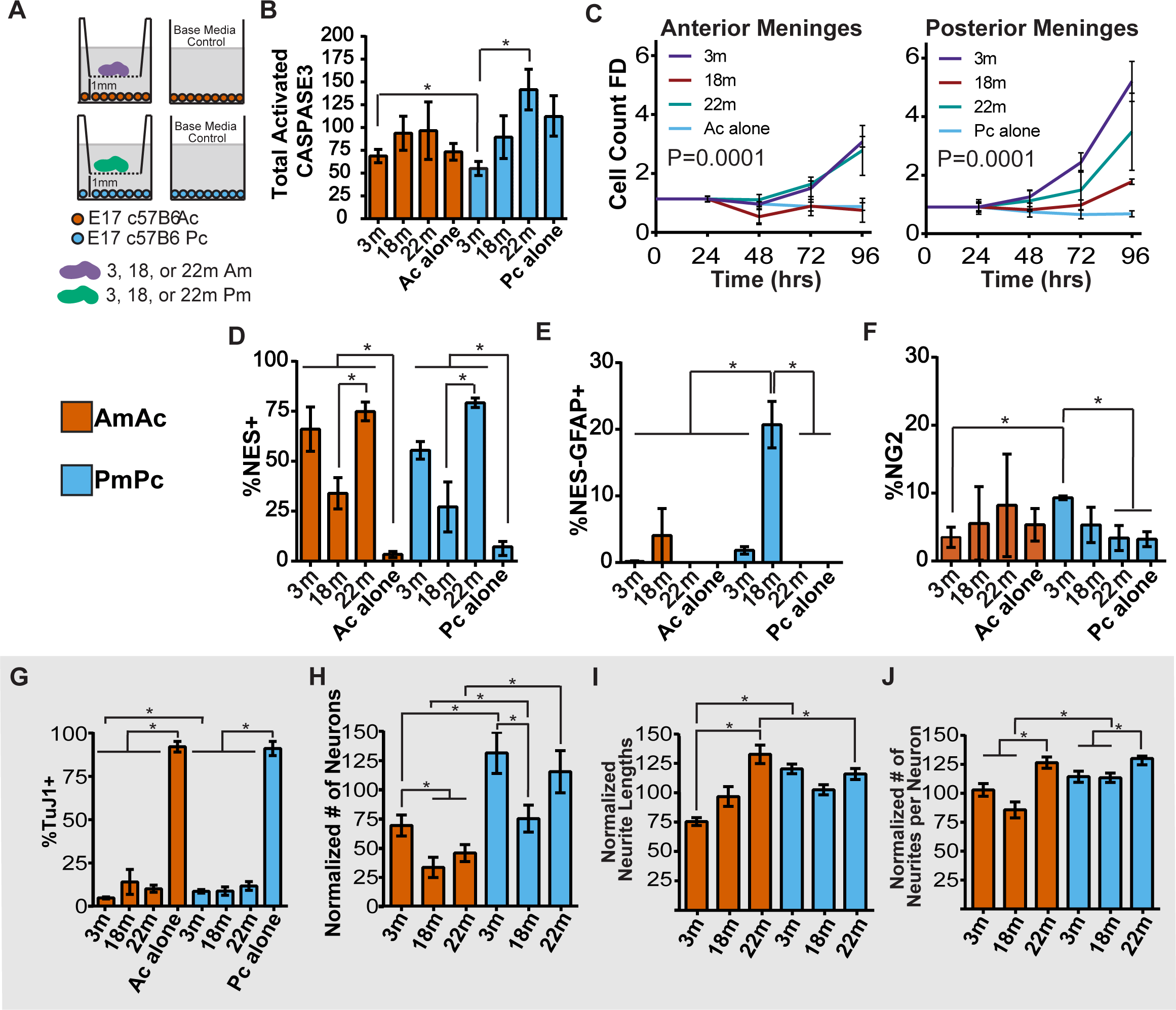
Leptomeningeal support of cortical cells changes with age in non-monotonic patterns. (A) Schematic of experimental design to assess the effect of co-culturing 3-, 18-, or 22-month- old leptomeninges with cortical cells: Am= anterior leptomeninges (purple); Ac= anterior cortical cells (orange); Pm= posterior leptomeninges (green); Pc= posterior cortical cells (light blue) (n=3 in each condition). Control: cultures grown without leptomeninges; (B) Number of Caspase3+ cells normalized to total number of DAPI+ nuclei per well, normalized values were compared by unpaired Student’s t-test; asterisk indicates p value <0.05; (C) Ac or Pc cell count in co-culture for 96-hours with 3-, 18-, or 22-month-old Am or Pm in homotypic combinations (n=3 in each condition). Cortical cells were counted and normalized to cell number for each condition at 24 hours. Nonlinear regression models (second order polynomial line) were compared to determine if one curve fit all conditions by extra-sum-of-squares F test; a p<0.05 has a preferred model of different curves fitting each condition; (D-K) Ac or Pc were cultured with 3- 18- or 22-month leptomeninges or without leptomeninges (Ac or Pc alone); (D) % of NES^+^ progenitor cells; (E) % of NES^-^GFAP^+^ cells; (F) % of NG2^+^ cells; (G) % of TuJ1 neurons; (H) The number of TuJ1^+^ cells normalized to the number of TuJ1^+^ cells in Ac and Pc alone control conditions; (I) Neurite length as % of the average neurite lengths in Ac and Pc control conditions; (J) Number of neurites per neuron as % of the average neurites per neuron in Ac and Pc alone control conditions. Unpaired Student’s t test was used to determine statistical significance, (*=p<0.05). Error bars represent mean ± SEM.

Anterior cortical neurons were significantly fewer with 18- or 22-month compared to 3-month leptomeninges co-culture. 18-month posterior leptomeningeal co-cultures had significantly fewer neurons compared to 3-month co-cultures, although the number rebounded at 22-months (Fig. 5G,H). Unexpectedly, 22-month old leptomeninges promoted increased neurite lengths and more neurites than 3-month old anterior leptomeninges (Fig. 4I, J). In summary, leptomeninges show significant changes in cortical cell support with age. At 18-months, the ability of leptomeninges to support neurons and stimulate progenitor proliferation is markedly reduced.

**Figure 5.**
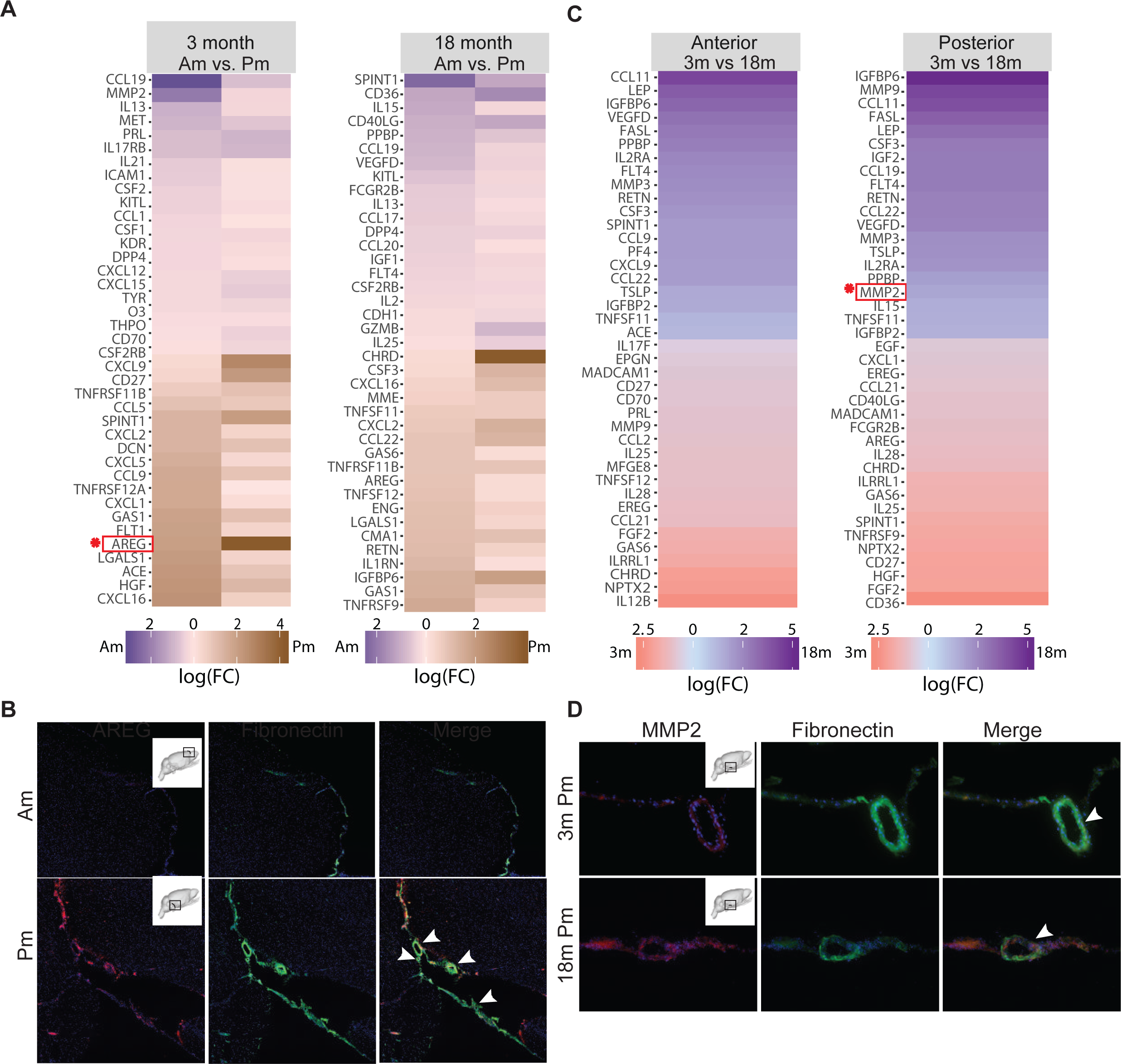
Secreted cytokine expression analysis identifies distinct protein enrichment profiles from 3-versus 18-month-old anterior (Am) and posterior (Pm) leptomeninges. (A) Heat map comparison of proteins secreted by 3- or 18-month-old anterior (purple) versus posterior (brown) leptomeninges that display differential or inverse expression when compared between 3- and 18-month-old leptomeninges (2 samples); (B) Immunostaining of AREG (red) in 3-month-old anterior and posterior leptomeninges. Fibronectin (green) immunostaining marks the leptomeningeal tissue and blood vessels; (C) Heat map comparison of cytokines secreted by anterior or posterior 3- (red) versus 18-month-old (blue) leptomeninges with differential or inverse protein expression between the anterior and posterior leptomeninges; (D) Immunostaining of MMP2 and Fibronectin in 3- and 18-month-old posterior leptomeninges. Heat maps represent log fold changes with at least a ≥1.5 fold difference in signal intensity considered significant.

### Regional and age differences in the leptomeningeal secretome

Given these significant effects, we wanted to examine the factors that could be providing support for the neural tissue. We first evaluated our snRNA-seq data by examining ligands expressed by each cell type (Table 1). Both endothelial and fibroblast populations expressed ligands that could promote growth and survival, such as Egf, BMPs, and Fgf14. We then evaluated the proteins released from anterior or posterior leptomeningeal regions at 3 and 18 months when the most profound functional differences were observed. Using a secreted protein array assay (G2000, RayBiotech), we compared enrichment of 144 proteins associated with cell survival, proliferation, differentiation, inflammation, and apoptosis. Proteins with the largest differences in 3-month anterior versus posterior and 18-month anterior versus posterior leptomeninges were identified (Tables S6-9). Regional and age-related differences in protein expression were determined using R.

We performed a pair-wise comparison of cytokines from anterior versus posterior leptomeninges dissected from individual animals (Table S6-10, Fig. 5A). For 3-month-old animals, proteins released at higher levels from anterior compared to posterior leptomeninges included CCL19, CCL1 and CXCL12 (chemotactic, immunoregulatory and inflammatory processes), and PRL, IL13, IL21, and CSF1 (growth-promoting). Those released more from posterior compared to anterior leptomeninges included CXCL9, CXCL16, and CXCL5 (chemotactic), HGF and AREG (growth-promoting), and CD27, LGALS1, and TNFRSF12a (apoptosis regulation). The top proteins released at higher levels from 18-month-old anterior compared to posterior leptomeninges include IL15, IL13, and CCL19 (immunoregulatory and inflammatory processes), IGF1 (insulin regulation), and FLT4 (growth factor binding and angiogenesis regulation). 18-month-old posterior leptomeninges secreted higher levels of CSF3, and AREG (growth-promoting proteins), IL1RN (growth factor binding), IGFBP6 and Resistin (RETN) (insulin regulation), and TNFSF11, TNFSF12, and GAS6 (apoptosis regulation). Supporting these findings, immunostaining showed higher levels of AREG in 3-month-old posterior compared to anterior leptomeninges (Fig. 5B).

To delve further into age-related differences, we examined the relative levels of cytokines released from leptomeninges at 3 versus 18 months (Fig. 5C). 3-month-old leptomeningeal conditioned medium was enriched for several anti-inflammatory (HGF,IL4, and IL10), proliferative (GAS6 and FGF2), and immune response-related factors (CXCL1, CD36, and IL17B) while 18-month-old conditioned medium was enriched for several growth-regulating and inflammatory-mediating proteins including LEP, IGFBP6, and IGFBP2, and TSLP, TNFSF11, MMP2, and IL15. By region, anterior leptomeninges with age release higher levels of proteins associated with the inflammatory response, immune cell activation, and TNF signaling including MMP3 and CCL11, while posterior leptomeninges in young animals release higher levels of proteins associated with cell migration or morphogenesis, such as GAS6, AREG, and EREG, which decline with age. Immunostaining for MMP2 in 3- and 18-month-old posterior brain sections confirmed higher expression in the aged leptomeninges (Fig. 5D). These findings demonstrate regional and age-related differences in the leptomeningeal secretome that may contribute to differences in paracrine effects on underlying neural tissue.

### Exploring a role for the meningeal secretome in schizophrenia

One of the unexpected findings from this study was the association of meningeal fibroblast and endothelial gene expression with mental illness, in particular schizophrenia. Schizophrenia typically has early adult onset, but it has been viewed as a progressive neurodegenerative disease that can worsen with age.^21^ With the demonstration that the regional leptomeninges release factors that impact cortical neuronal survival and morphology and the numbers of astrocytes and oligodendrocytes, we hypothesized that changes in these factors could contribute to the pathogenesis of schizophrenia. To explore this possibility, we downloaded bulk RNA-seq data from the BrainSeq Phase 2 project^22^ (https://eqtl.brainseq.org/phase2/) taken from the dorsolateral prefrontal cortex (DLPFC) of schizophrenic and control patients. This dataset is comprised of 199 male and 101 female control samples and 104 male and 49 female schizophrenia samples. The incidence of schizophrenia is similar between males and females, although females typically have a later adult onset.^23^

We first identified differentially expressed (DEX) genes between control and disease within each sex, i.e. control females vs schizophrenic females, control males vs schizophrenic males, using a Wilcoxon-rank test. After identifying DEX genes from the schizophrenia samples, we utilized a ligand-receptor database^24^ to identify receptors in the patients that had ligands expressed by the meningeal cells in a region- or age-dependent manner in our snRNA-seq data (Fig. 6A-B). We first tested to determine if DEX receptors in the schizophrenia data were enriched for association with leptomeninges secreted ligands and we found a high enrichment for both the female (p= 1.584996e-27) and male (1.153724e-25) datasets. We observed about a third more potential interactions between the meningeal ligands and the receptors of the DLPFC of female schizophrenic patients over males. For example, the ligand *AVP* (Arginine Vasopressin) secreted by *Fibro1* cells in the young posterior leptomeninges along with the DLPFC receptors *Avpr1a* (Arginine Vasopressin Receptor 1A) and *Oxtr* (Oxytocin Receptor) has previously been implicated in schizophrenia^25^ and this network was present in the female but not the male data. Several subtle changes exist between the female and male networks that include different receptors for the same ligands. *Il13ra2* and *Tlr2* receptors, both interaction partners of the IL4 (Interleukin 4) ligand that is secreted more by the young posterior leptomeninges, are shown to be differentially expressed in the female schizophrenic patients but not males. On the other hand, *Il4ra* expression is lower in male patients but unchanged in females. The IL13RA2 receptor, which is increased in female schizophrenia patients, is thought to alter *Il4* signaling and promote TGFB production.^26^ Illustrating a potential consequence of the changes in *Il13ra2* expression, a TGFB allele associated with increased risk of schizophrenia in female but not male patients,^27^ has been demonstrated to increase TGFB secretion.^28^ *Il4ra,* which is downregulated in male patients, promotes IL4 signaling, and decreases in IL4 levels have been implicated in early onset schizophrenia, which is more common in males.^29-31^ Overall, these results indicate that it will be worthwhile to consider the role that leptomeninges-derived factors may play in the development of schizophrenia and other mental disorders.

**Figure 6.**
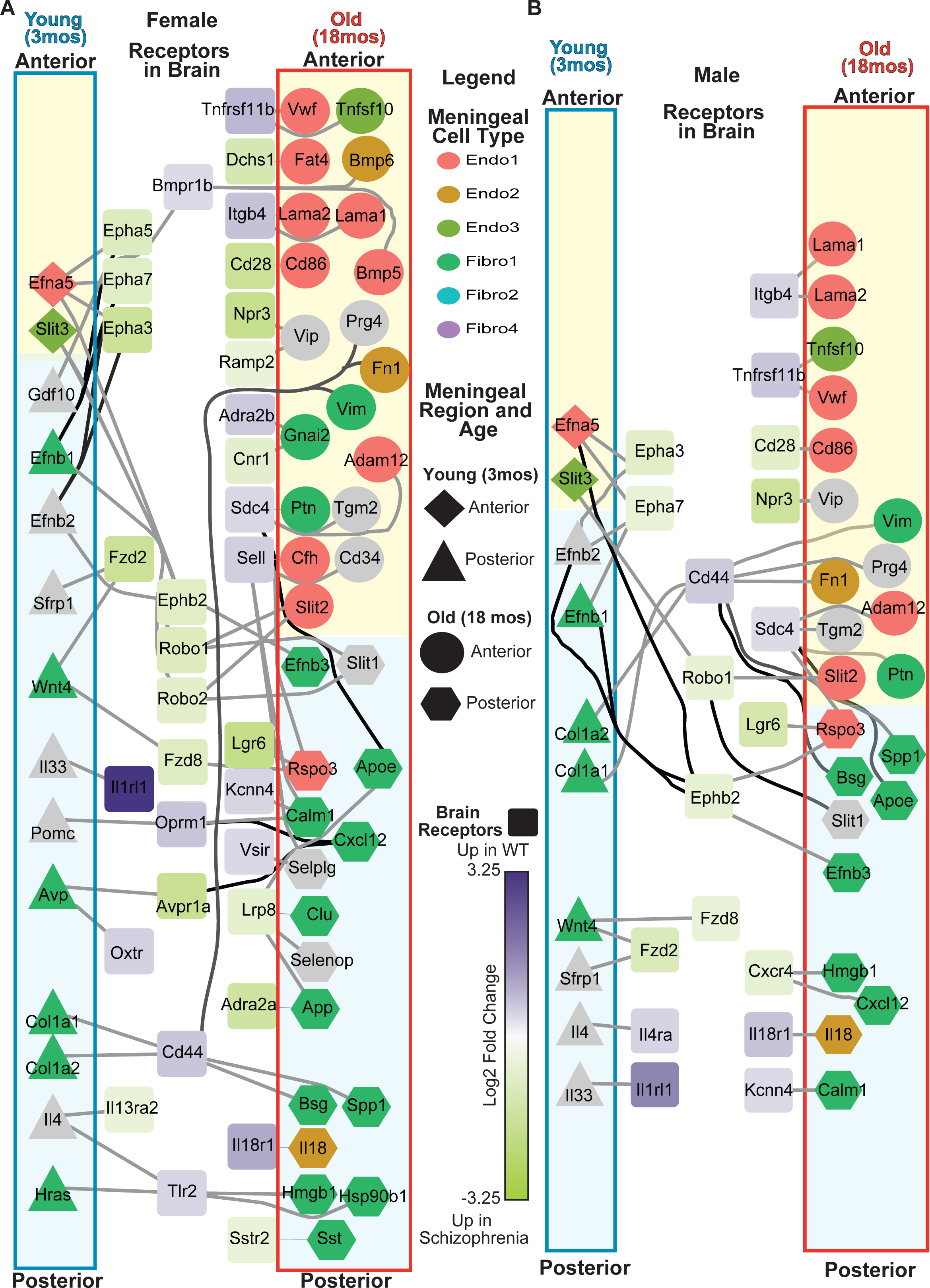
Uncovering potential crosstalk across age and region between the leptomeninges ligands and receptors in the brain in schizophrenia patients. Schizophrenia expression data was downloaded from BrainSeq phase II project (https://eqtl.brainseq.org/phase2/) and ligand receptor pairs were downloaded from the CellTalkDB (http://tcm.zju.edu.cn/celltalkdb/download.php). Ligands were identified as significantly different in leptomeningeal subpopulations, or with region or with age and intersected with DEX receptors from (A) female and (B) male schizophrenia patients. The color of a ligand identifies the subpopulation (grey= no subpopulation) and the shape of the ligand identifies the age and region with which it is associated. The receptors are colored based on the expression level in schizophrenia patients compared to controls with purple being high in controls and green high in schizophrenia.

## Discussion

Adult leptomeninges support underlying neural tissue, protect against injury, promote tissue repair, and maintain a homeostatic environment. Much is known about the global benefit of leptomeninges, but the concept that adult leptomeninges are regionally specialized remains largely unexplored. Because leptomeningeal cells penetrate the brain in the perivascular spaces, they are in close proximity to brain cells. Using functional assays, snucRNA-seq, and assays of protein release, we revealed that adult leptomeninges exhibit regional patterning with differential ability to support cortical neurons, neurite outgrowth, glia and glial progenitor cells. Moreover, we demonstrated that meningeal function changes with age, affecting the ability of each region to support underlying neural tissue. Our findings expand upon prior pioneering studies showing that young adult leptomeninges stimulate radial-glial like progenitor cells at the pial surface^6,17,18^ to encompass meningeal stimulation of other neural cells, and by identifying regional and age-related differences in adult meningeal-progenitor cell interactions. Hence, this study opens avenues of exploration to define the impact of regionalized, specialized support provided by the leptomeninges to underlying CNS regions, and the consequences of impairment or dysfunction in this relationship.

Our snRNA-seq data of meningeal cells demonstrated a strong regional identity that was a primary source of variance in the dataset. We identified several distinct aspects of meningeal- cortical cell functional interactions that differed with meningeal region, the most profound effect being a dramatic mitogenic stimulus of glial cells. The adult 3-month anterior leptomeninges stimulated production of anterior cortical GFAP^+^NES^+^ cells, most likely cortical astrocyte precursors, while 3-month posterior leptomeninges stimulated oligodendroglia production from posterior cortical progenitors. Further, we found differential effects of leptomeningeal co-culture on numbers of cortical neurons of different subtypes, neurite production and growth. Deficits in neurite outgrowth and the genes regulating this process have been associated with bipolar disorder ^32^ and schizophrenia,^33^ which may connect our transcriptomic disease-association analysis and functional results. Anterior leptomeninges supported CTIP2+ subcortical projection neurons greater than posterior, while the reverse was the case for SATB2+ callosal projection neurons. Hence impairments in these relationships could conceivably have differential effects on cortical neuron populations that are differentially vulnerable to particular diseases. We found that with age, leptomeningeal support of cortical neurons and glia changes significantly. Both anterior and posterior leptomeninges exhibited reduced ability to support cortical cell survival and growth with age, with several regionally distinct differences. For example, aged posterior leptomeningeal-derived factors caused increased cortical cell apoptosis gene expression and reduced oligodendroglial expansion compared to 3-month-old posterior leptomeninges, and we noted that aged posterior leptomeninges gene expression showed enrichment for programmed cell death terms. We suggest that such changes could contribute to differences in cell survival, myelin dysfunction and remyelinating repair seen with aging and associated with mental disorders such as depression, bipolar disorder, and schizophrenia.^34-36^

Following these analyses, we examined the proteins secreted from leptomeninges and found definitive evidence that in young 3-month-old adults, different proteins are released from the leptomeninges from different brain regions. For example, prolactin, PRL, is released more from 3-month anterior versus posterior leptomeninges. PRL stimulates astrocyte proliferation and expression of differentiation markers^37^, which could contribute to the pro-astrogliogenic effect of anterior leptomeninges we observed. Amphiregulin (AREG), a mitogen for astrocytes and fibroblasts that supports progenitor cell expansion and is neuroprotective, is enriched in 3-month-old posterior leptomeninges compared to anterior.^38-40^ AREG enrichment may explain the enhanced ability of posterior leptomeninges to stimulate cortical progenitor cell expansion. 3-month posterior leptomeninges also expressed higher levels of hepatocyte growth factor (HGF) which is known to stimulate oligodendrocyte proliferation ^41^ and might explain their pro-oligodendroglial effects that we observed in co-cultures.

With age, several factors associated with inflammation, mental disorders, and neurodegenerative disease show increased release from the leptomeninges, including MMP2, FLT4, CSF3, and IL15.^42-46^ These findings corroborate prior studies that link meningeal-derived inflammatory factors to progression of neurodegenerative disease and mental disorders.^1,47^ Our findings that secreted factors from the leptomeninges can interact with brain receptors altered in schizophrenic patients strengthen the potential link between meningeal-derived environmental effects and schizophrenia progression. One of the ligands showing increased expression in the posterior leptomeninges with age in the *Endo2* population, *Il18*, has been associated with decreased cognition in schizophrenia patients.^48^ MMP2, another molecule linked with schizophrenia^46^ and Alzheimer’s disease (AD),^49^ can cleave CXCL12, which is expressed by the leptomeninges, into a truncated neurotoxic peptide that induces apoptosis and neurodegeneration.^50,51^ MMP2 is abundantly expressed in aged leptomeninges (Fig. 4H) and if it actively cleaves CXCL12 in the posterior leptomeninges this may help explain the increased apoptosis observed in our experiments. An increase in both CXCL12 and MMP2 in the leptomeninges, if occurring in humans, could contribute to age-related changes including inflammation and neurodegeneration seen in schizophrenia and AD.^42,52^ From our network analysis examining potential meningeal factors influencing schizophrenia pathogenesis, we highlighted IL13RA2 and we found by protein array analysis that IL13 secretion is increased in the anterior leptomeninges, providing another potential mechanism influencing the etiology of schizophrenia.

In conclusion, leptomeninges are an essential mediator of brain homeostasis and further investigation of regional differences in cell composition, paracrine function and impact on neural cells will provide valuable insight into their contributions to different regions of the CNS in health, normal aging and with neurodegenerative diseases.

## Acknowledgements.

We thank Steve Lotz and other members of the NSCI NeuraCell core facility for ICELL8 system support, and Meryl Lindsay and Brian Unruh for technical assistance. This study was supported by the National Institute of Neurological Disorders and Stroke (NINDS) of the National Institutes of Health award R35NS097277 (ST).

## Author contributions

CA performed experiments, data analysis, and wrote the first draft of the manuscript.

SG assisted with experimental design and dissection.

ML assisted with experimental design.

TK supported the single nuclei analysis.

FF performed single nuclei isolation, library prep, and mapping of single cell data

YW performed the tissue isolation and assisted in single nuclei isolation

NB performed single nuclei and protein array analysis, performed data analysis, and wrote the manuscript.

ST conceived the topic, helped with data analysis and interpretation, and wrote the manuscript.

## ONLINE METHODS

## EXPERIMENTAL MODEL AND SUBJECT DETAILS

All handling procedures were reviewed and approved by the University at Albany IACUC committee. Timed pregnant C57BL/6 female mice were purchased from Taconic Biosciences and 3-, 18-, and 22-month-old male C57BL/6 mice from the National Institute on Aging. Mice had no history of previous experimentation. All mice were housed in a humidity and temperature-controlled environment under a 12:12 light/dark cycle with a standard free-range diet. Upon arrival, mice were given at least 48 hours to acclimate to our animal facility before experimentation. All mice were visually inspected and observed to confirm good health. To minimize batch effects, age-matched mice from different cohorts were used for experimental replicates.

### Cortical tissue dissociation and dissociation

Timed pregnant C57BL/6 mice were anesthetized with isofluorane and embryos removed. Embryo ages were verified using Theiler Springer-Verlag, 1989 anatomical guidelines.^53^ Only embryos at 17 days of gestation were used for co-culture experiments. Anterior and posterior cerebral cortex tissue was dissected in hibernation medium ^54^. Cortical tissue pooled from 4-5 embryos was enzymatically dissociated at 37°C for 45 minutes with light trituration every 15 minutes in high glucose (4.5 mg/mL) DMEM (Life Technologies) or DMEM without phenol red (Life Technologies) with papain (20U/mL, Worthington), sodium pyruvate (1mM, ThermoFisher Scientific), N-Acetyl-L-cysteine (1mM, Sigma-Aldrich), L-glutamine (2mM, ThermoFisher Scientific), and DNase (0.024 mg/mL, Sigma-Aldrich). Cortical cells were washed 3X with high glucose DMEM (Life Technologies). Single cells were obtained by filtering dissociate with a 40μm cell strainer (Falcon), then staining with 0.08% trypan blue (Corning) to quantify live versus dead cells, counted using a hemocytometer under 10X magnification (Zeiss Axiovert microscope).

### Meningeal tissue dissection and dissociation

C57BL/6 male mice were anesthetized with isofluorane. 6 mice were pooled for each experimental replicate. Brains were submerged in hibernation medium and anterior and posterior leptomeninges dissected (Fig. S1). Tissue was dissociated at 37°C for 10 minutes with light trituration every 5 minutes in Hanks Balanced Salt Solution (HBSS) (ThermoFisher Scientific) with trypsin (0.25%, ThermoFisher Scientific) and DNase (0.024 mg/mL, Sigma-Aldrich). Meningeal cells were washed 1X with high glucose DMEM (Life Technologies). Red blood cells were lysed with red blood cell lysis buffer (Biolegend) for 10 minutes on ice, followed by 3X washes with high glucose DMEM (Life Technologies). Single cells were obtained by filtering with a 40 μm cell strainer (Falcon), then stained with 0.08% trypan blue (Corning) to identify live versus dead cells counted using a hemocytometer under 10X magnification (Zeiss Axiovert 40C).

### Cell culture

Anterior and posterior cortical cells were plated onto poly-L-ornithine (Sigma-Aldrich) coated 24-well plates at a density of 25,000 cells per well in high glucose DMEM (Life Technologies) with sodium pyruvate (1mM, ThermoFisher

Scientific), N-Acetyl-L-cysteine (1mM, Sigma-Aldrich), L-glutamine (2mM, ThermoFisher Scientific), N2 supplement (1mM, ThermoFisher Scientific Sci.), and B27 Supplement (ThermoFisher Scientific); culture medium was supplemented with FGF2 (10 ng/mL ThermoFisher Scientific) for the first 24 hours. 1 hour after cell plating, a transwell insert (Corning) containing 10 mg of anterior or posterior meningeal tissue was inserted into the well: Young 3-month-old anterior or posterior leptomeninges were co-cultured with anterior or posterior cortical cells in homo- or heterotypic combinations; Aged 18- or 22 month-old anterior or posterior leptomeninges were plated with anterior or posterior E17 cortical cells in homotypic combinations. Equal weights of anterior or posterior leptomeningeal tissue, confirmed by measuring total protein using a BCA assay (Fig. S1B), were placed into each transwell compartment above cerebral cortical cells plated in equal cell numbers per well (Fig. 3A). Every 24 hours, cultures were fed by exchanging half the medium with fresh medium. After 96 hours, cortical cultures were fixed with 4% paraformaldehyde for 20 minutes followed by 3X washes with sterile PBS (Gibco Life Technologies) and stored at 4 °C until further processing.

### Immunocytochemistry

Cells were fixed with 4% paraformaldehyde (Santa Cruz) for 20 minutes at room temperature, then washed 3X with PBS. Cells were multiplex stained sequentially, as described below, washing 1X after incubating with each primary antibody and 3X after incubation with secondary antibodies:

Cells were first stained with NG2 primary antibody in PBS for 1 hour at room temperature (1:200, Millipore/Chemicon) followed by a 1hour incubation with goat anti-rabbit IgG Cy3-546 secondary antibody in PBS (1:800, Invitrogen).Cells were then blocked for 1 hour with 2% bovine serum albumin (Sigma-Aldrich), 10% normal goat serum (Vector Laboratories), and 0.2% PBS with 0.05% Triton X (Sigma-Aldrich). Cells were stained overnight with SATB2 primary antibody (1:25, Abcam), then incubated for 1 hour at room temperature with goat anti-mouse IgG1-488 secondary antibody (1:500, Invitrogen). Cells were then stained with β-tubulin III primary antibody for 45 minutes at room temperature (1:500, Sigma-Aldrich) then goat anti mouse IgG2b Cy5-647 secondary (1:300, Invitrogen). Finally, cells were incubated with the nuclear dye DAPI for 5 minutes (1:1000, Invitrogen). Cell staining results were manually counted from digital pictures taken using an inverted epifluorescence microscope at 10X magnification (Zeiss). Cells were then stained with primary antibodies to: Nestin (1:10, Developmental Studies Hybridoma Bank), GFAP (1:200, Dako), and CTIP2 (1:200, Abcam) for 1 hour at room temperature followed by a 1 hour incubation at room temperature with IgG1-488, IgG Cy3-546, and IgG1-488 secondary antibodies, respectively. Multiplexed stained cells were imaged at the same X,Y vernier reading as the first round of digital imaging and then manually counted. All manual counting was completed using AxioVision software (Zeiss).

Cell death was measured by quantifying activated Caspase-3 levels in cortical cells following co-culture with young or old leptomeninges. Cortical cells were fixed and blocked as previously described and then stained with primary antibody to activated Caspase-3 (1:300, Cell Signaling) overnight at 4 °C followed by a 1-hour incubation with IgG Cy3-546 secondary antibody. A total of 300 10X images were taken of each experimental condition using a plate reader (Cytation5 Imaging reader). Staining was counted using Cell Profiler software (version 3.0.0, Broad Institute).

### Total protein analysis

A Coomassie (Bradford) protein assay was used to assess total secreted protein accumulation every 24 hours in culture media with or without leptomeninges. Total protein was measured by referencing a standard of serially diluted bovine serum albumin (following manufacturer’s instructions). All samples were diluted to remain within the acceptable reference concentration range. Total protein concentrations of standards and samples were measured with a Nanodrop instrument (Thermo Scientific).

Relative secreted protein was determined following 4 days of leptomeningeal culture either alone or co-cultured with cortical cells using a secreted protein array (G2000, RayBiotech). Methods to process conditioned media followed manufacturer’s guidelines.

Relative secreted protein expression levels, median protein signal intensities, and background subtractions were configured for each experimental condition (Raybiotech). Samples with proteins having at least a ≥1.5 fold difference in signal intensity were considered significantly different (Table S6-S9). Automatic computation for all sample comparisons were configured using R (code available at https://github.com/neural-stem-cell-institute/Meninges).

### Single cell separation, cDNA preparation, RNA Sequencing, mapping, and analysis

Anterior and posterior leptomeninges tissues were isolated from six young (3 months) and six old (18 months) mice. Tissue lysis and nuclei isolation were performed according to the 10x genomics nuclei isolation for single cell RNA sequencing protocol (CG000124, Rev D), without myelin removal. Nuclei were stained with Syto64 (5 µM) for 15 minutes and single cell library preparation was performed using the SMART-Seq ICELL8 Application Kit (Takarabio, catalog no. 640221) according to the manufacturer’s instructions. RNA sequencing was performed at Genewiz using NovaSeq6000 Illumina platform. Read counts were demultiplexed and mapped to mouse genome GRCm38 using annotation mm10 with Cogent NGS analysis pipeline (Takarabio, V1).

Data was analyzed using R software with Bioconductor packages (code available on github: https://github.com/neural-stem-cell-institute/Leptomeninges. Cells having greater than 1000 features and library sizes of at least 30,000 reads were included for analysis. Genes with greater than a ratio (exon+intron counts/ intron counts) of 100 were excluded from analysis. Raw FASTQ files and a .RDS of the Seurat object are available for download from GEO (GSE242810).

## QUANTIFICATION AND STATISTICAL ANALYSIS

GraphPad Prism version 5.0 was used for statistical analysis. All biological replicates (n), and significance parameters are described in the figure legends. Biological replicates consisted of independent experiments run sequentially. To account for possible batch effects, age-matched mice from different cohorts were used for each experimental replicate. All graphed data are shown as a mean ± the standard error of the mean (SEM). Comparisons of normally distributed data were tested using a two-sided, unpaired Student’s t-test. N.S. p > 0.05, *p< 0.05. Data values comparing growth over time were compared with nonlinear regression models fit to a second order polynomial line. The nonlinear regression models determined if one curve fit all conditions by extra-sum-of-squares F test. Total protein was normalized by computing z-scores for aging comparisons (3 or 18-month old Am or Ac cultured alone and 3 or 18-month old Pm or Pc cultured alone), and standard scores were then used to normalize data values over time in hours. Protein levels at 96 hours were tested with a Welsh’s ANOVA. Protein array heat maps represented a significance threshold defined as having a log fold change with ≥1.5 fold difference in signal intensity.

## DATA AND CODE AVAILABILITY

Raw reads were submitted to GEO (GSE242810). All R code written for this analysis is available at https://github.com/neural-stem-cell-institute/Meninges.

---

**Supplemental Figure S1.**
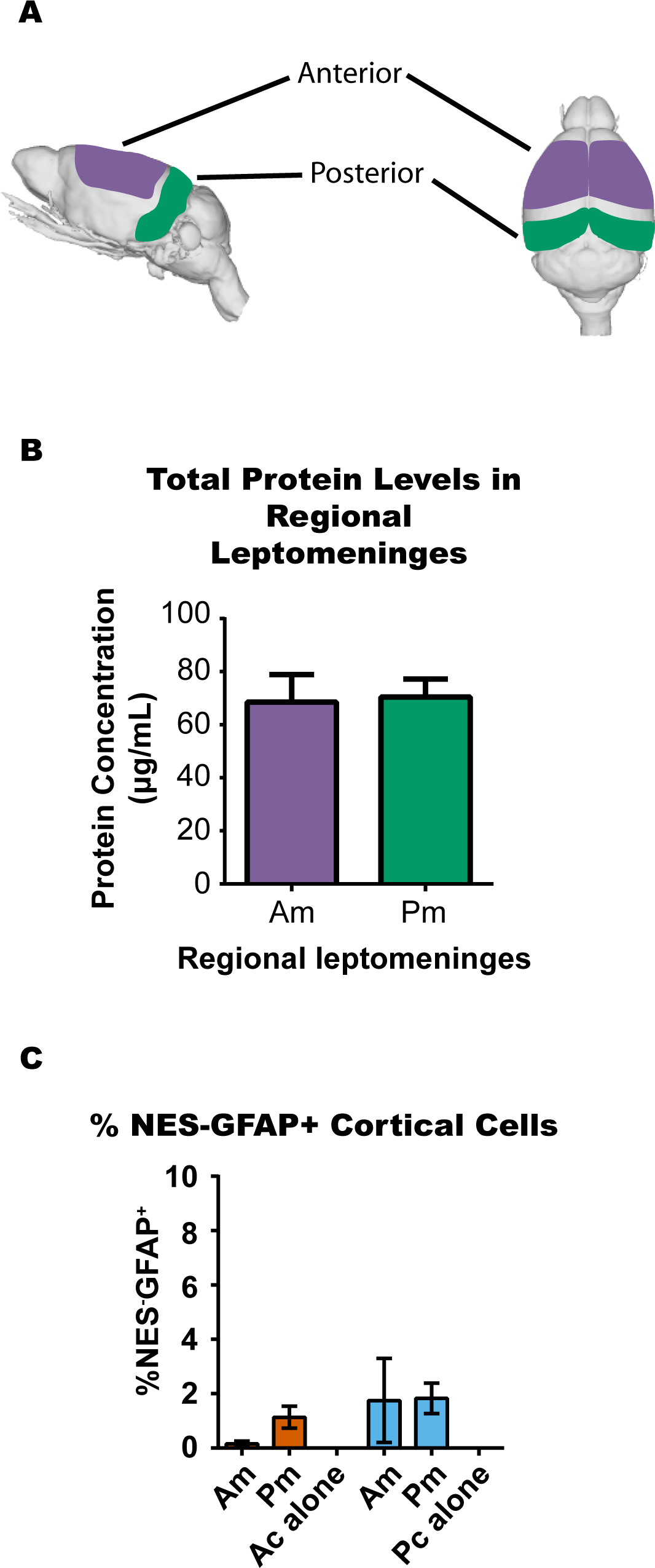
(A) Diagram of showing the locations of ‘Anterior’ and ‘Posterior’ leptomeninges as defined throughout this study. (B) Protein levels were measured by BCA assay. (C) Cultures were stained for NESTIN and GFAP. Shown are the NESTIN negative and GFAP postiive cells, which are mature astrocytes.

**Supplemental Figure S2.**
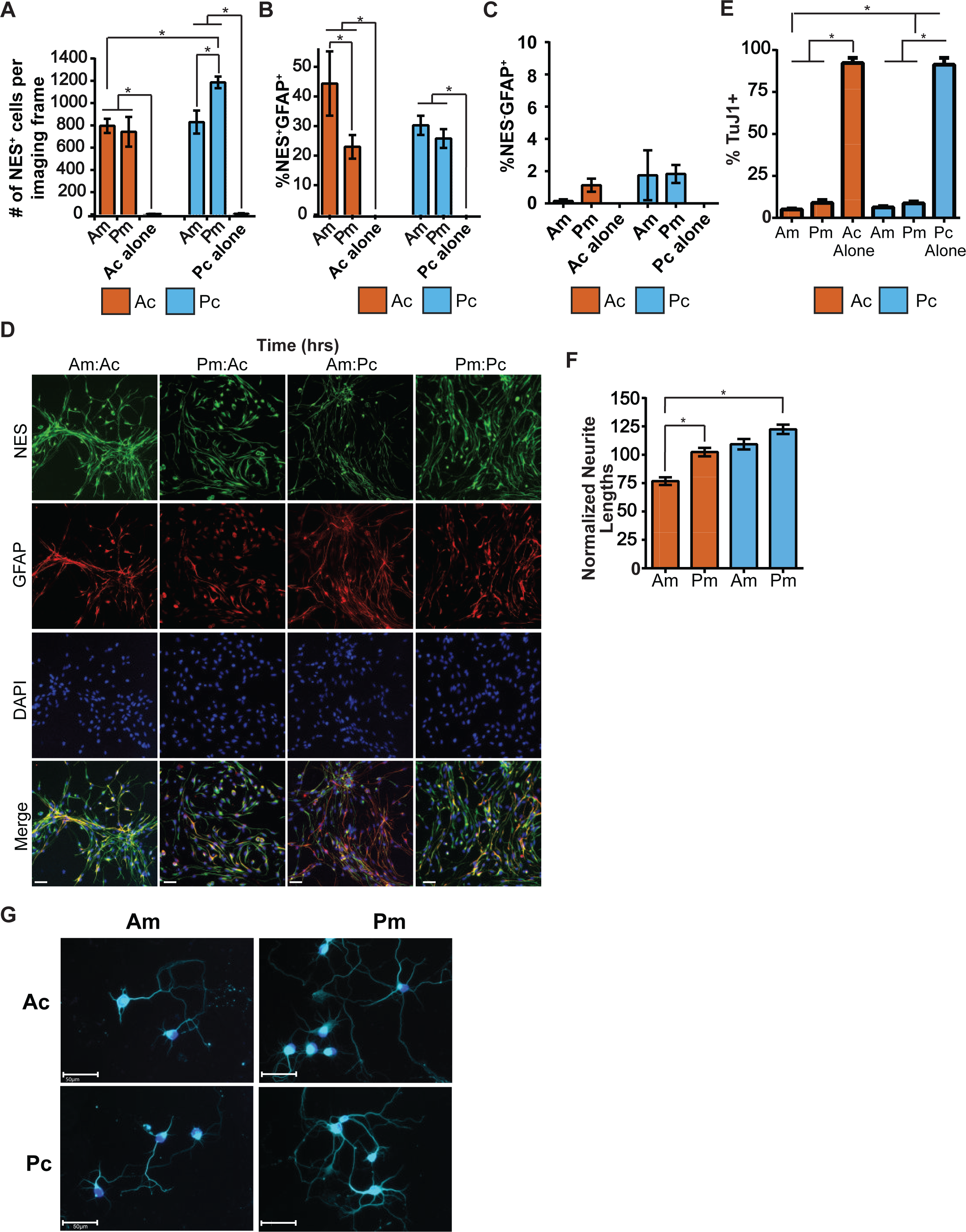
(A) Total number and NES+ progenitor cells; (B) % of NES+ GFAP+ or (C) NESGFAP+ cells; (D) immunofluorescence images illustrating differences in NES+ and GFAP+ cells in Ac and Pc co-cultured with Am or Pm (Scale bar=100μm) (E) % TuJ1+ cells (neurons) in each condition; (F) Normalized neurite lengths in Am and Pm co-culture over the average neurite lengths in Ac or Pc alone control conditions; (G) Immunofluorescence images illustrating differences in neuron morphology after TuJ1 (b-tubulin III) staining (scale bar= 50um); F)

